# Accurate sub-population detection and mapping across single cell experiments with PopCorn

**DOI:** 10.1101/485979

**Authors:** Yijie Wang, Jan Hoinka, Teresa M Przytycka

## Abstract

The identification of sub-populations of cells present in a sample, and the comparison of such sub-populations across samples are among the most frequently performed analyzes of single cell data. Current tools for these kind of data however fall short in their ability to adequately perform these tasks. We introduce a novel method, PopCorn (single cell sub-<u>Pop</u>ulations <u>Co</u>mpariso<u>n</u>), allowing for the identification of sub-populations of cells present within individual experiments while simultaneously performing sub-populations mapping across these experiments. PopCorn utilizes several novel algorithmic solutions enabling the execution of these tasks with unprecedented precision. As such, PopCorn provides a much needed tool for comparative analysis of populations of single-cells.

## 1 Introduction

Recent technological advances have facilitated unprecedented opportunities for studying biological systems at single-cell level resolution. For example, single-cell RNA sequencing (scRNA-seq) enables the measurement of transcriptomic information of thousands of individual cells in one experiment. Analyses of such data provide information that was not accessible using bulk sequencing which can only assess average properties of cell populations. Single cell measurements however can capture the heterogeneity of a population of cells. In particular, single cell studies allow for the identification of novel cell types, states, and dynamics [1, 2, 3, 4]. The benefits of single cell data however come at the cost of unique computational challenges [5]. These challenges emerge from the stochasticity of single cell experimental data as well as from the multitude of questions that are being addressed with this technology.

One of the most prominent uses of the scRNA-seq technology is the identification of sub-populations of cells present in a sample and comparing such sub-populations across samples [6, 7, 8, 9, 10, 11, 12, 13]. Such information is crucial for understanding the heterogeneity of cells in a sample and for comparative analyses of samples from different conditions, tissues, and species. While some information about sub-population structure can be gained from data visualization methods such as the dimensionality reduction technique tSNE [14], relaying on visualization approaches alone can be highly misleading. To address this challenge, Butler at al. developed a computational approach enabling the identification of single cell populations across data sets [15]. This method is based on the Correlated Components Analysis (CCA) followed by an alignment of the CCA basis vectors between the data sets. The method has been implemented as a part of the popular software package Seurat [15]. While Seurat’s approach provides the first step towards addressing this important problem, however, it focuses on aligning cell populations across experiments ignoring that there exist specific clustering structures in individual experiments. Although the aligned data can then be clustered to reveal sub-populations and their correspondence, solving the sub-population mapping problem by performing global alignment first and followed by a clustering method overlooks the original information about sub-populations existing in each experiment and an approach that addresses this problem directly might be more successful. To address this critical gap we developed a new approach, single cell sub-Populations Comparison (PopCorn) that allows for comparative analysis of two or more single cell populations.

There are two key ideas behind PopCorn that are fundamental for the accuracy of our approach. The first idea is to identify sub-populations of cells present within individual experiments simultaneously with performing sub-populations mapping across these experiments rather than identifying the sub-populations first and mapping them later. This allows for integrating information across experiments thus reducing noise. The second key innovation consists of a new approach to identify sub-populations of cells within a given experiment. Unlike simple clustering approaches, PopCorn utilizes Personalized PageRank vectors [16] and a quality measure of cohesiveness of a cell population (introduced in Supplementary Materials A) to construct an auxiliary sub-population co-membership propensity graph to guide the process of identifying such sub-populations.

We tested the performance of PopCorn in two distinct settings. First, we demonstrated its potential in identifying and aligning sub-populations from single cell data from human and mouse pancreatic singe cell data [15]. Next, we applied PopCorn to the task of aligning biological replicates of mouse kidney single cell data [17]. PopCorn achieved a striking improvement over the existing tool.

Consequently, and as a result of our integrative approach, PopCorn provides novel and unmatched tool for comparative analysis of single-cells populations.

## 2 Method

Informally, a sub-population of cells should include cells that have a common expression pattern (consistency) which are distinct from the expression patterns of other cells (separation). However, applying this principle in the context of single cell experiments is non-trivial. Given the stochastic nature of single cell experiments, some sub-populations can be well separated in one experiment whereas the separation can be less pronounced in another - either due to technical issues or due to biological differences between the samples. In addition, the stochasticity of the experiment introduces noise to the readout of the expression level of individual genes in individual cells, which might impact the accuracy of the assessment of the similarities between cells.

The main idea of the PopCorn is based on the simultaneous identification of sub-populations of cells present within individual experiments with performing sub-populations mapping across experiments. To this purpose, PopCorn integrates two objective functions aimed at (i) ensuring a meaningful partition of cells into sub-populations within each individual experiment and (ii) ensuring the consistency of these partitions across the experiments. To jointly optimize these two objectives, we construct two weighed graphs. The first graph, also known as sub-population co-membership propensity graph, encodes the propensity of cells belonging to the same sub-population for any two cells form the same experiment (*A* in Fig. 1) (we use same label to denote a graph and its matrix representation). The criterion used for constructing this graph is key to the efficiency of our method and is described in the next subsection and Supplementary Materials A. The second graph, a multipartite graph, encodes pairwise similarities of cells from different experiments (*B* in Fig. 1). Following the construction of these two graphs, the final step of the method consists in solving a *k*-partition problem that simultaneously takes into account constraints encoded by both graphs.

**Fig. 1:**
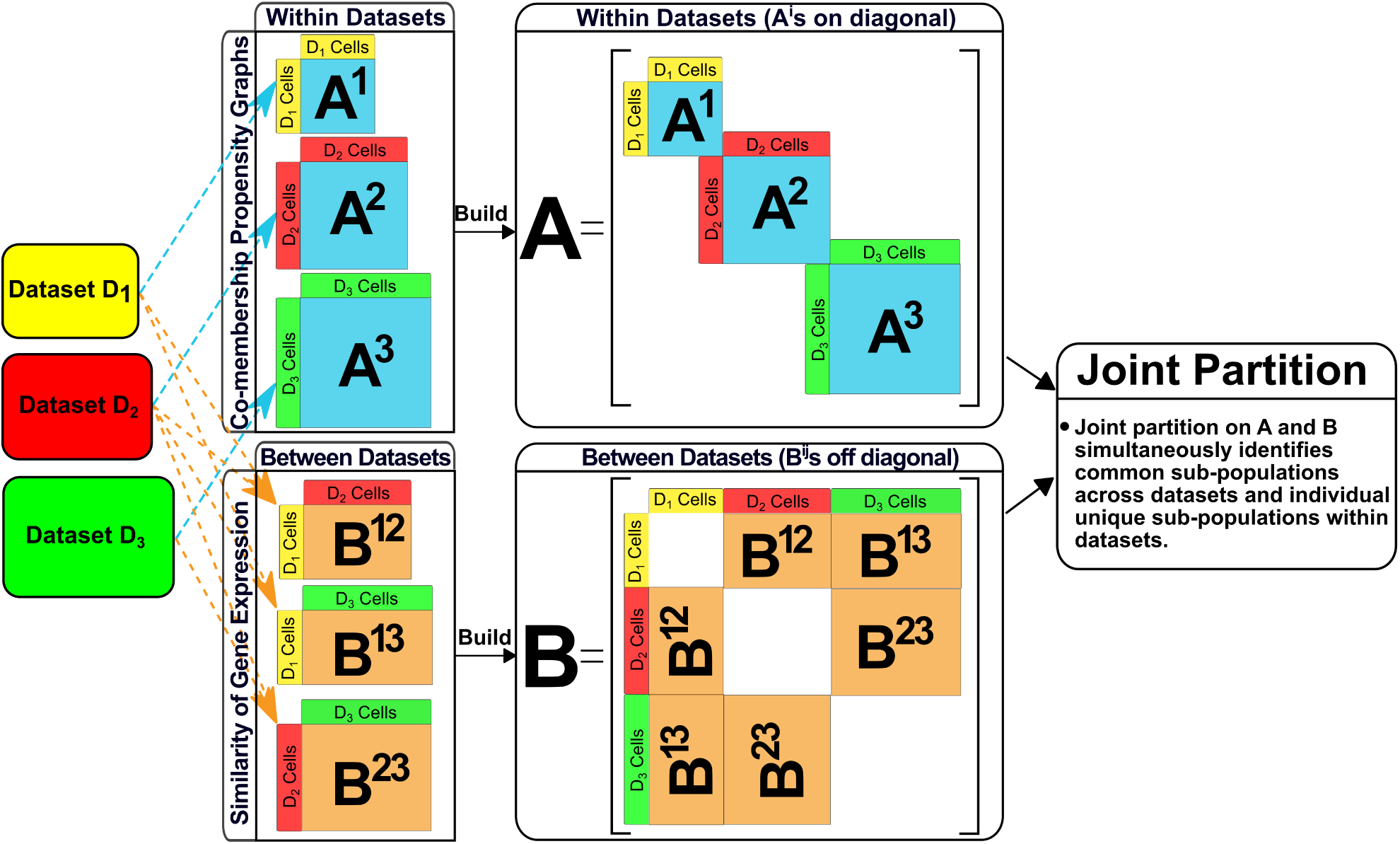
The work flow of PopCorn. For each experiment *i* we first construct, matrix *A*^*i*^ representing the propensity of each pairs of cells to be in the same sub-population (this is achieved by using personalized PageRank method, see Section 2.1 and Supplementary Materials A). Matrix *A* summarizes all matrices *A*^*i*^ and is constructed by placing these on the diagonal of *A*. Next, we construct matrices *B*^*i, j*^ which encode similarities between pairs of cells form different experiments *i* and *j* (see Section 2.2). *B* is constructed by placing the similarity matrices for all pairs of experiments off-diagonal as illustrated in the figure. We then perform joint partition of the graphs represented by matrices *A* and *B* by applying semi-definite programming to solve the problem.

In the subsequent sections, we outline the main ideas behind our approach while more technical details are deferred to the supplementary materials. We additionally note that the subsequent sections assume the existence of *q* scRNA-seq data sets denoted by *D*_1_, *D*_2_, *…, D*_*q*_ where a data set *D*_*i*_ covers *N*_*i*_ genes over *M*_*i*_ cells.

### 2.1 Construction of the sub-population co-membership propensity graph

The objective of sub-population identification is to partition cells into groups while optimizing for consistency within each group and separation between the groups.

To address these challenge, we compute *sub-population co-membership propensity graph, A*^*i*^, for every experiment *D*_*i*_. *A*^*i*^ consists of a weighted graph with nodes corresponding to the cells from experiment *D*_*i*_ and edge weight represents the propensity of a given pair of cells to be in one cluster (one sub-population). To estimate such propensity, each cell “votes” which other cells should be put in the same sub-population with itself. The voting process utilizes a personalized PageRank vector on the expression similarity graph (see (5) and Supplementary Materials A for a precise definition) for the cells in experiment *D*_*i*_. For any cell, the ranking of the cells in its personalized PageRank vector utilizes a measure of expression consistency and a separation to ensure the desired properties for the sub-populations proposed (See Supplementary Materials A for more details). An edge (*l, m*) is included in the graph sub-population co-membership propensity graph *A*^*i*^ if this cells *l* and *m* obtained at least one vote to be in the same sub-population. The weight of each edge equals to the number of votes received by the given pair. Note that partitioning *A*^*i*^ into *k* sub-graphs provides a method to uncover the population structure in experiment *D*_*i*_ which is of an independent interest. However if performed in isolation, such sub-population assignment would not benefit from the information contained in the data from other experiments. Thus graphs *A*^*i*^ are used jointly with the information represented by graph *B* that encodes pairwise similarities between cells from different experiments as described below.

### 2.2 Solving joint sub-population identification and mapping problem

Solving the joint sub-population identification and mapping problem translates into grouping the cells into a set of clusters such that the resulting partition is optimized for grouping cells of similar expression pattern within data sets while ensuring cells of the same kind are also aligned across data sets.

To this end, we encode complementary information regarding the sub-population co-membership propensity and the pairwise cell-similarities from different experiments in two graphs *A* and *B* respectively.

*A* corresponds to the union of all graphs *A*^*i*^ as defined in the previous section. Let *N* = Σ_*i*_ *N*_*i*_ be the overall set of cells, arranged such that cells from the same data set have consecutive indices. *A ∈* ℝ^*N×N*^ is then constructed by assembling the adjacency matrices *A*^*i*^, *i* = 1, 2, *…, q* into a block-diagonal matrix, i.e. 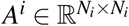 are arranged along the diagonal as shown in Fig. 1.

*B* in turn consists of a *q*-partite graph recording the similarity between cells across different experiments based on gene expression. Formally

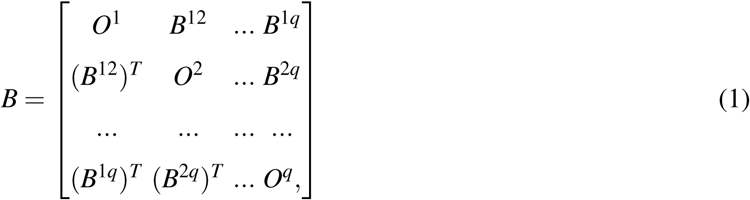

where 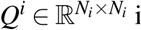 is an all zero matrix and 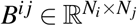 measures expression consistency between cell from different experiments (see definition in (6).) Note that *A, B ∈* ℝ ^*N×N*^ have the same dimension.

Given these two adjacency matrices, we then compute a partition of the cells into *k* clusters that respects the connectivity defined by both graphs. To accomplish this, we first normalize both matrices as described in Supplementary Materials B.1 and define the normalized Laplacian matrix of *A* and *B* as *L*_*A*_ and *L*_*B*_ respectively. To encode the assignment of cells to a sub-population we define an assignment matrix *Y*_*N×k*_, where *N* is the total number of cells, as follows:

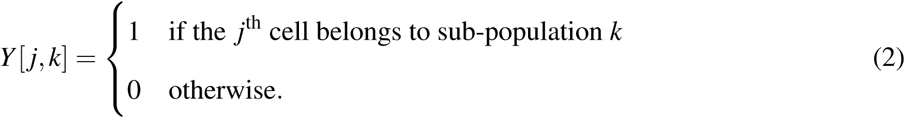

Note that given the normalized Laplacian matrix *L*_*X*_, where *X* is either *A* or *B*, the problem of finding the optimal partition into *k* sub-populations that respects connectivity defined by matrix *X* is equivalent to the normalized *k*-cut problem [18, 19] and can be expressed as

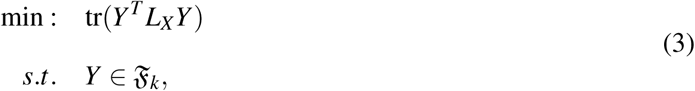

where 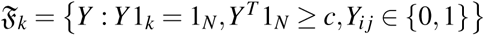.

In particular, if *X* = *A*, this leads to clustering the scRNA-seq data sets into *k* different sub-populations, based solely on experiment specific features (similarity expression pattern between different cells in the same experiment) without using any information from other experiments. In contrast if *X* = *B* we will find the best alignment between the cells across experiments ignoring sub-population structure withing each experiment.

Accordingly, a *k*-partition formulation respecting both matrices can be defined as

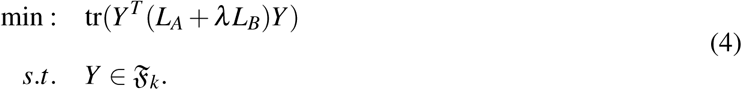

were the parameter *λ* defines a scalar weight relation between the two sets of edges. To find the optimal solution we use a semi-definite programming (SDP) relaxation approach as described in the Supplementary Materials B.

### 2.3 Expression similarity between cells

Expression similarity is defined differently for cells form the same experiment compared to cells from separate experiments, although both share a first common step: the identification of highly variable genes (HVGs) [20] *Ω*^*i*^ in each data set *D*^*i*^ and consequent utilization the normalized expression data 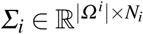 of those HVGs to compute expression consistency.

For cells *l* and *m* belonging to the *i*th data set *D*^*i*^, their expression consistency 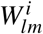 is computed using the cosine similarity as follows:

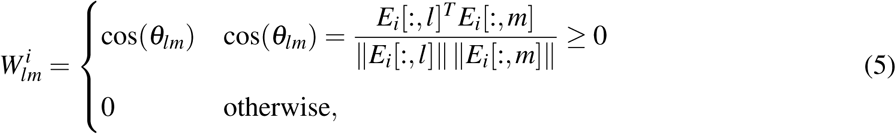

Here, 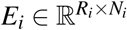 is derived by applying Principle Component Analysis (PCA) to *Σ*_*i*_ where *R*_*i*_ corresponds to the number of principle components.

To compute expression consistency between cells across different experiments, the posibility of different scRNA-seq data sets having different sets of highly variable genes (HVGs) needs to be taken into account. To account for this variability, we compute the similarity between cells *l* and *m* across data sets using coexpressed HVGs only:

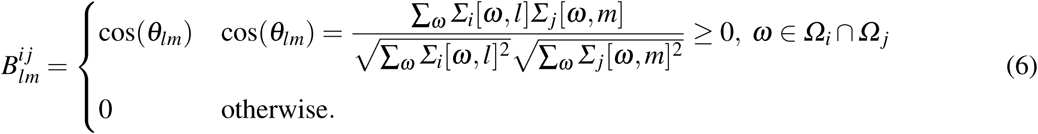

## 3 Results

### 3.1 Comparative analysis of single-cell RNA-seq experiments across species

To demonstrate the capabilities of our approach, we first applied PopCorn on two scRNA-seq data sets from different species and compare its performance to methods that first integrate and align the data and then cluster the integrated data. In particular, we obtained both, human and mouse pancreatic cell transcriptomes from GEO Series accession number GSE84133. The human scRNA-seq data set contains 8,629 cells from 13 cell types whereas the mouse scRNA-seq data set includes 1,886 cells from 11 cell types, 10 of which are shared by both data sets. In addition, the human scRNA-seq data set has 3 individual cell types that do no appear in mouse scRNA-seq data set. We performed a comparative analysis on both data sets to identify *individual sub-populations* which only contain cells from a single data set, and *common sub-populations* which include cells from more than one data set. We further utilize the labels provided in [15] as our gold standard to validate the performance of the comparative analysis. The results generated by our PopCorn approach are then benchmarked against the results of methods that first integrate and align the data and then cluster the integrated data. The comparison between PopCorn and Seurat alignment [15] method followed by Louvain clustering algorithm [21] (Seurat + Louvain) is illustrated in the following paragraphs and the comparison between PopCorn and a recently proposed method Scanorama [22] followed by Louvain clustering algorithm [21] (Scanorama + Louvain) is reported in Supplementary Materials D.2. For a detailed description of the parameter selection regarding both methods, we refer the reader to Supplementary Materials D.1.

In order to ensure an impartial comparison, we define several metric scores which evaluate the results of the competing methods from distinct, yet comlementary contexts. The first metric, *R*_*ccsp*_, aims at measuring how accurate a method can group cells of the same cell type across data sets together and is defined as the ratio of the number of identified *corresponding common sub-populations* to the total number of identified *common sub-populations*. Next, we evaluate how many common cell types across data sets and individual cell types can be recovered by a method via *R*_*ucct*_ (the ratio of uncovered common cell types to the total number of common cell types) and *R*_*uict*_ (the ratio of uncovered individual cell types to the total number of individual cell types). Last but not least, we utilize *Accc* and *Acci* to quantify the purity of the identified *common sub-populations* and *individual sub-populations*. Purity in this context refers to the percentage of the majority cell population in an identified sub-population. The detailed definitions of all metric scores can be found in Supplementary Materials C.

Fig. 2 A illustrates two t-distributed stochastic neighbor embedding (tSNE) plots that summarize the results generated by PopCorn for the two benchmark data sets. Although the layouts of the cell sub-populations in human and mouse are distinctive in appearance, the figure illustrates PopCorn’s ability to identify *common sub-population* (sub-population #1 to #10) and *individual sub-populations* (sub-population #11 to #13). Fig. 2 B visualizes the content of each identified sub-population discovered by PopCorn and the Seurat alignment method. The right figure indicates that for sub-population # 9 and sub-population # 10, Seurat assigned cells of different types in human and mouse into one group. Specifically, sub-population # 9 classified human acinar cells and mouse beta cells as one common sub-population. Human acinar cells however are a unique sub-population that does not have a correspondence population in mouse experiments. In contrast, our PopCorn method successfully identified the human acinar cells as a unique sub-population (sub-population # 12 of our in the left figure of Fig. 2). In addition, we observed that sub-population # 10 of Seurat contains both mouse activated stellate and quiescent stellate cells and human activated stellate and quiescent stellate cells, but fails to designate them into separate sub-populations. PopCorn however correctly assigned mouse activated stellate and human activated stellate cells into one sub-population (subpopulation # 9 of our PopCorn), and mouse quiescent stellate and human quiescent stellate cells into another sub-population (sub-population # 10 of our PopCorn). Last but not least, Seurat fails to identify any known *individual sub-populations*. However, PopCorn identifies 2 known *individual sub-populations* (acinar and T-cell cells). Fig. 2 C compares the results of PopCorn and Seurat on the metrics as defined above and shows that PopCorn outperforms Seurat on all metrics scores.

**Fig. 2:**
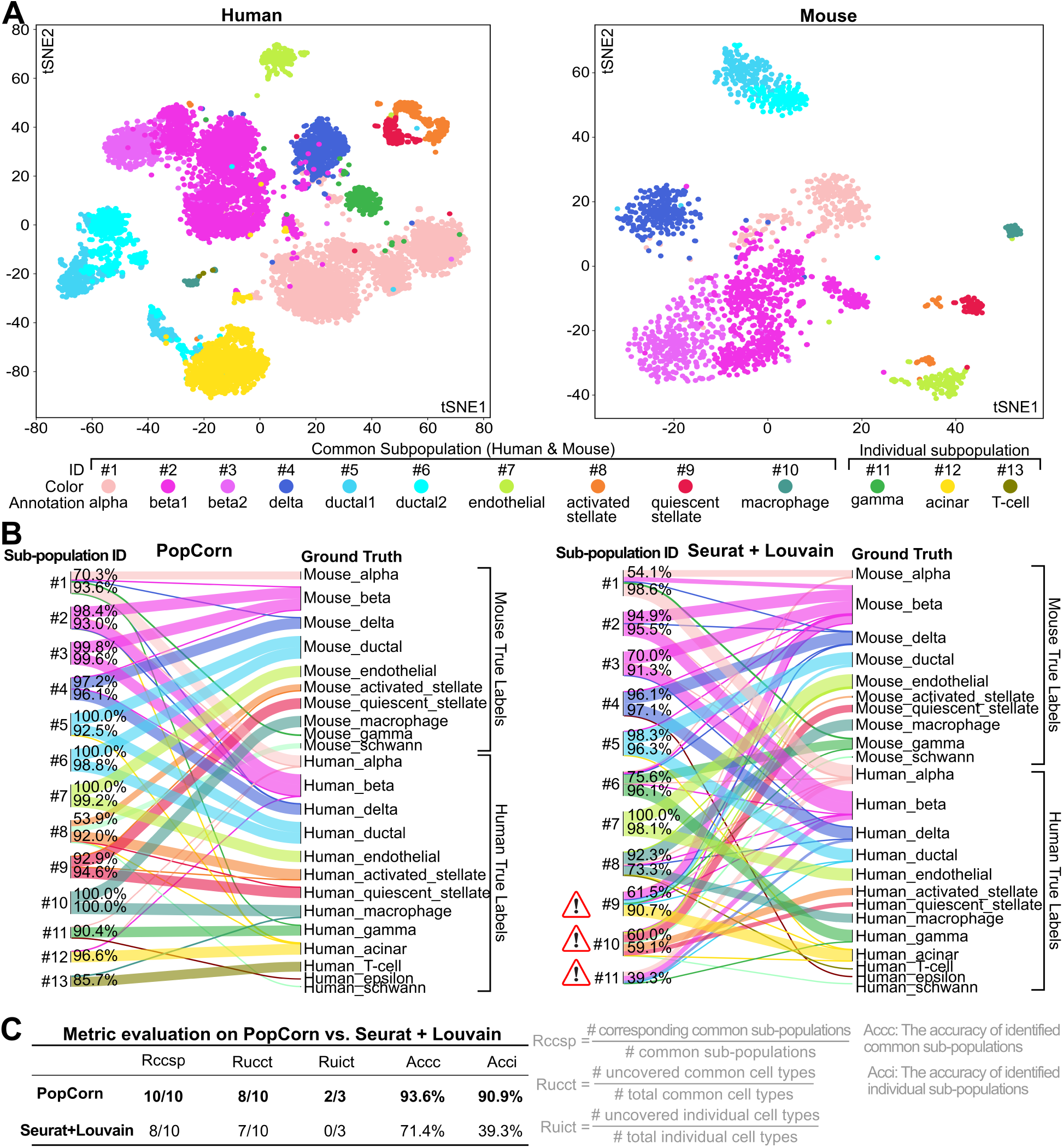
(A) Two t-SNE plots for human and mouse scRNA-seq data sets, respectively. Different colors indicate different cell annotations, which can be determined via true labels (Supplementary Materials C). Cells of the same color denote a sub-population identified by PopCorn. (B) Sankey diagrams of the resulting mapping between identified sub-populations to ground truth labels of both PopCorn and Seurat. The width of the flow bar is proportional to the *Acc*_*ssp*_ score (the accuracy of a *split sub-population*) as defined in Supplementary Materials C.2. The *Acc*_*ssp*_ scores for the majority of the cell types in each identified sub-population are given at the beginning of the flow bar. As shown, all *common sub-populations* (sub-population #1 to #10) identified by PopCorn are *corresponding common sub-populations*. In addition, PopCorn identifies three *individual sub-populations* (sub-population #11, #12, and #13) that are annotated to acinar, gamma, and T-cell, and acinar and T-cell are known to be unique cell populations in the human experiments. In contrast, sub-population #9 of Seurat + Louvain incorrectly assigned acinar cells in human and beta cells in mouse into a single population; sub-population #10 of Seurat + Louvain designated both activated stellate and quiescent stellate cells in human and mouse into one population but failed to separate them. In addition, Seurat + Louvain failed to identify any individual cell populations. #11 contains only cells from human but is a mixture of different cell types. (C) Comparison of PopCorn and Seurat on various metric scores that are defined in Supplementary Materials C.

### 3.2 Comparative analysis of single-cell RNA-seq data sets with multiple replicas

Next, we tested the robustness of PopCorn to perform under the presence of multiple replicas, a crucial biological imperative to monitor the quality and repeatability of an experiment. Computational approaches processing the resulting data are expected to be resilient against reasonable technological and biological variations and to produce consistent results across replicas. To test this, we applied PopCorn and Seurat + Louvain to mouse kidney scRNA-seq data recently published in [17] and performed comparative analysis on the four replicas. The replicas, identified by GEO association numbers GSM2871706, GSM2871707, GSM2871708, and GSM2871709, contained 2,943 cells, 5,060 cells, 1,383 cells, and 2,704 cells, respectively. Moreover, the four replicas have 16 cell types in common and no distinctive individual cell types. Finally, and analogous to the previous analysis procedure, the cell labels provided in [17] were used as ground truth to evaluate the performance of each method. The comparison on the same data sets between PopCorn and Scanorama + Louvain is shown in Supplementary Materials D.3. The parameter selection of both methods can be found in Supplementary Materials D.1.

Fig. 3 summarizes the findings of applying the above described test scenario on PopCorn and Seurat and shows that PopCorn outperforms Seurat on all metric scores. As shown in Fig. 3 A, PopCorn successfully identifies 18 common sub-populations, all of which are *corresponding common sub-populations*. In contrast, Seurat identifies 19 *common sub-populations*, 5 of which are *non-corresponding common sub-populations* erroneously grouping together cells of different types in different replicas. Specifically, sub-population #15 of Seurat incorrectly assigns DCT cells in replica 3 into a group of CD-PC cells from replica 1, 2, and 4 and sub-population #16 groups Macro cells in replica 2 with Fib cells in replicas 3 and 4. In addition, sub-population #17 contains B-lymph cells in replica 1 and Endo cells in replica 2 while sub-population #18 falsely assigns T-lymph cells in replica 3 and NK cells in replicas 1 and 4 together, and sub-population #19 mistakenly groups Macro cells in replica 1 together with B-lymph cells in replicas 2, 3, and 4. It is also worth noting that out of the 16 common cell types as stated in [17], PopCorn identified a total of 11 as evidenced by the number of unique annotations for *common sub-populations*. In contrast, Seurat was only able to uncover 6 common cell types (Fig. 3 A, right). This prompted us to further investigate the properties of the remaining common cell types that were not uncovered by PopCorn. Interestingly, we found these cell types, Podo, CD-Trans, Novel1, Neutro and Novel2, to have particularly small cell populations of 10, 56, 42, 25, and 10 cells respectively corresponding to 0.1, 0.5, 0.4, 0.2, and 0.1 percent of all cells in 4 replicas.

**Fig. 3:**
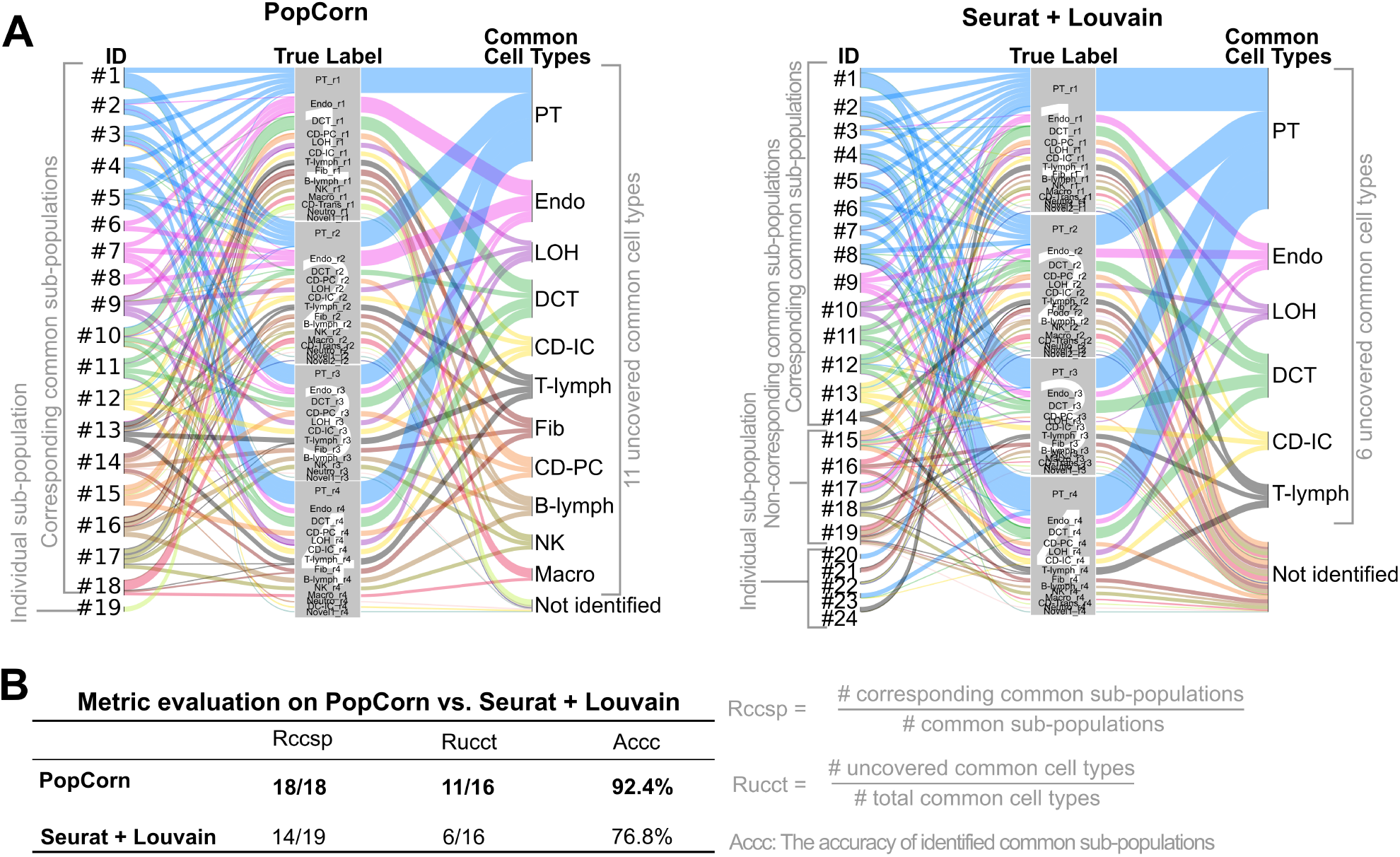
(A) Sankey diagrams of the results mapping between identified sub-populations to ground turth labels of both PopCorn and Seurat. The numbers in the background of the center gray area correspond to the respective replica indexes. The width of the flow bar is proportional to the *Acc*_*ssp*_ score (accuracy of a *split sub-population*) defined in Supplementary Materials C.2. The *Acc*_*ssp*_ scores are given in Supplementary Materials D.3. As shown, all *common sub-populations* identified by PopCorn are *corresponding common sub-populations*. In contrast, 5 out of 19 common sub-populations identified by Seurat are *non-corresponding common sub-populations*, which assinged cell types of different kinds in individual data sets together. Furthermore, PopCorn uncovered 11 common cell types whereas Seurat uncovered only 6. (B) Comparison between PopCorn and Seurat on mouse kidney single cell data sets. PopCorn outperforms Seurat on all metrics as described in Supplementary Materials C.

## 4 Conclusions

We developed PopCorn, a new method for the identification of sub-populations of cells present within individual single cell experiments and mapping of these sub-populations across the experiments. In contrast to alternative approaches PopCorn performs these two tasks simultaneously by optimizing a function that combines both objectives. When applied to complex biological data the results produced by our approach are of unprecedented quality, robustness, and reproducibility across replicas that was not available in previous method. Several innovations developed in this work contributed to this success. First, incorporating the above mentioned tasks into a single problem statement was crucial for integrating the signal form different experiments while identifying sub-populations within each experiment. Next, the sub-population co-membership propensity graph introduced here to guide sub-population identification in individual experiments significantly aids the reliable identification of groups of similar cells that are well separated from the remaining populations. Taken together, these two ideas enable highly accurate identification of sub-populations and superior alignment of cells across populations.

With these qualities, PopCorn has great potential to become a fundamental tool in the analysis of single cell data.

## 5 Availability

A preliminary reference implementation of PopCorn is available upon request.

## 6 Acknowledgements

This research was supported by the Intramural Research Programs of the National Library of Medicine.

## Supplementary Materials

### A Construction of co-membership propensity graphs

In this section, we introduce how we construct a co-membership propensity graph based on a similarity matrix. Let us assuming we have a similarity matrix *W* that is derived by using (5). *W* can be viewed as the weighted adjacency matrix for a weighted un-directed graph *G*(*V, E*). *W*_*i* *j*_ encodes the similarity between node (cell) *i* and node (cell) *j*.

For a given node *v ∈ V*, the group 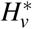 of nodes that tend to be in the same partition with *v* can be found by using the personalized PageRank vector of cell *v* [16]. The personalized PageRank vector *p*(*α, v*) of *v* on *G* is the stationary distribution of the random walk on *G*, in which at every step, the random walker has the probability of *α* to restart the random walk at *v* and otherwise performs a lazy random walk. Mathematically, *p*(*α, v*) is the unique solution to

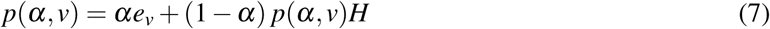

where *α ∈* [0, 1] is the “teleportation” constant, *e*_*v*_ is the indicator vector of *v* (where *e*_*s*_ = 1 if *s* = *v* and *e*_*s*_ = 0 if *s ≠ v*) and 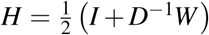 is the underlying probability transition matrix of the lazy random walk. *D* is a diagonal matrix with the weight sum of each node *d*(*v*) = Σ_*m*_ *W*_*vm*_, *∀v ∈ V* on its diagonal *D*_*vv*_ = *d*(*v*). We apply the modified local algorithm in [16] to efficiently approximate 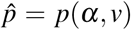. The algorithm in [16] is for unweighted graphs and we extend the algorithm for weighted graph as shown in **ApproximatePageRank weight(***v*, *α*, *ε*), which unities **Push**_*u*_(*p*, *r*).

**Push**_*u*_(*p*, *r*)

1. Let *p*′= *p* and *r*′= *r*, except for the following changes:

a. *p*′(*u*) = *p*(*u*)+ *αr*(*u*)
b. 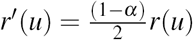
c. For each *v* such that 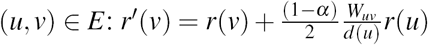
2. Return (*p*′, *r*′)

Then we sort the nodes based on 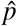 and attain an ordered set *H*_*v*_ = {*h*_1_, *h*_2_, *…, h*_*n*_}, *h*_*i*_ *∈V*, whose elements satisfy 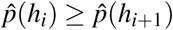. It is easy to verify that when *α >* 0.5, so that node *v* is always on top of the list *H*_*v*_, meaning *h*_1_ = *v*. Therefore, we use *α* = 0.8 through out the paper to make sure that *v* is in 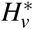. Based on the ordering in *H*_*v*_, we generate a collection of sets *S* _*j*_ = {*h*_1_, *h*_2_, *…, h* _*j*_ for *j ∈* {0, 1, *…, |H*_*v*_*|*}, which we call *sweep sets* [16]. We let

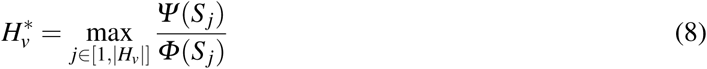

**ApproximatePageRank weight(***v*, *α, ε*)

1. Let *p* = 0 and *r* = *e*_*v*_.
2. While 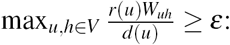
  a. Choose any node *u* where 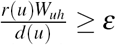
  b. Apply **Push**_*u*_ at node *u*, and update *p* and *r*
3. Return *p* with 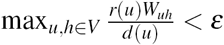

be the node set including *v* that has propensity to be well-separated (characterized by *Φ* (*·*)) and densely connected (characterize by *Ψ* (*·*)). *Ψ* (*S*) is the weighted density between nodes in set *S* defining as

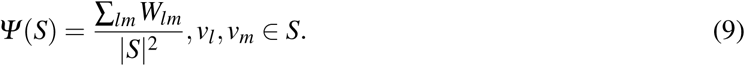

And *Φ* (*S*) is the conductance of set *S* that characterizes separation of *S*.

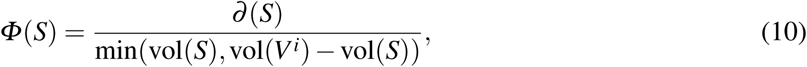

where *∂* (*S*) = Σ_*i* *j*_ *W*_*i* *j*_, *i ∈ S, j ∉ S* and vol(*S*) = Σ_*x∈S*_ *d*(*x*). 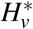 can be represented by an indicator vector 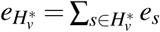. After obtain 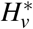 for every *v ∈V*^*i*^, we can compute the adjacency matrix of the co-membership propensity graph for *G* as follows

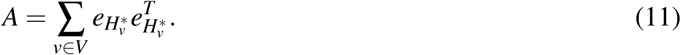

Based on the method introduced above, we can compute *A*^*i*^ for each single cell data set *D*^*i*^.

### B Optimization

#### B.1 Normalized Laplacian matrix

For adjacency matrix *A*, the normalized Laplcian matrix of *A* is defined as 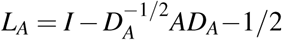, where *I* is an identify matrix and *D*_*A*_ is the weighted degree matrix with *D*_*ii*_ = Σ _*j*_ *A*_*i* *j*_ on its diagonal. Similarly, the the normalized Laplcian matrix of *B* is 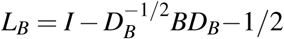.

#### B.2 SDP relaxation of (4)

The joint partition problem (4) is a NP-hard problem. In order to efficiently obtain promising results, we relax the problem into SDP relaxation. Based on previous results [19], we know the following problems (P1) and (P2) can be relaxed to (SDP1) and (SDP2).

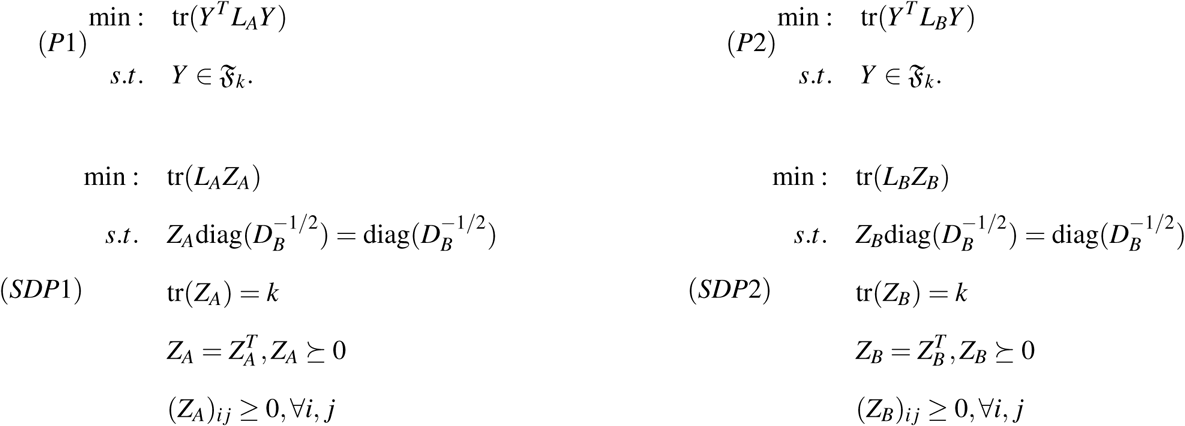

In those SDP relaxations 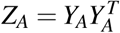 and 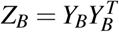, where 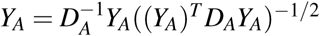 and 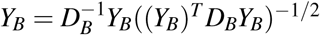. Comparing (SDP1) and (SDP2), we find that they have exact the same constraints except the first ones. We therefore relax the first constraint of (SDP1) and let *Z* = *Z*_*A*_ = *Z*_*B*_ and combine (SDP1) and (SDP2) to obtain our final SDP formulation as follow.

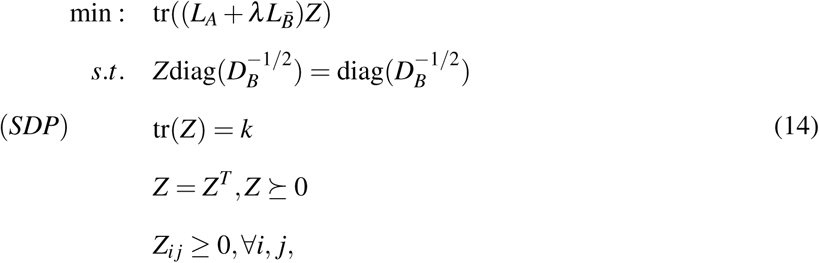

Problem (14) can be solved by well-established toolbox and we use cvxpy to solve the relaxed SDP problem.

#### B.3 Rounding

In general, due to relaxation, the optimal solution of (SDP) is not feasible for (4). Therefore, we need to recover a closest feasible solution to the original problem (4). We treat row *i* of *Z*, the optimal solution of (SDP), as the feature of cell *i*. Then we apply k-means to *Z* 100 times and pick one k-means solution which yields the minimum objective function value of (4).

#### B.4 Preclustering

Since our objective function unitizes a full *N × N* matrix 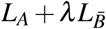, the memory usage of the algorithm may be prohibitive for increasing number of data sets. This motivates us to use *supercells*, which can be obtained by over-segmentation of the data sets (in our case, hierarchical clustering is used but any other superpixel identification methods in the image processing filed can be applied). Using *N*_*S*_ supercells is equivalent to constraint *Z* to be block-constant and thus reduces the size of the SDP to a problem of *N*_*S*_ *× N*_*S*_.

### C Evaluation metric

#### C.1 Terminology

Given *q* data sets {*D*^1^, *D*^2^,*…, D*^*q*^}, we use *C* = {*C*^1^,*C*^2^,*…,C*^*q*^} to present a sub-population that is identified by a method, where *C*^*i*^ is a *split sub-population* that only contains cells from data set *D*^*i*^. We call a sub-population a *individual sub-population* once *C* only contains cells from one data set (*C*^*i*^ *≠* Ø and *C*^*j*^ = Ø, *∀ j ≠ i*). When *C* contains cells from more than just one data set, we call *C* a *common sub-population*.

#### C.2 C.2 Annotation of a *split sub-population*

For a non-empty *split sub-population* 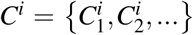 in *C*, we can annotate each cell 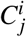 in *C*^*i*^ by its ground truth label 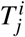 We use 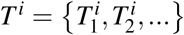 to present the label set of *C*^*i*^, where 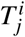 is the ground truth label for 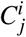. Assuming we have *L* ground truth label *T* = {*T*_1_, *T*_2_, *…, T*_*L*_}. Then the annotation 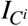 of the *split sub-population C*^*i*^ can be found by

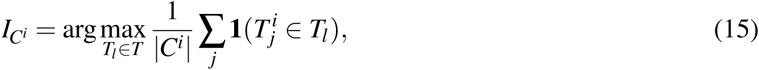

where **1**(*·*) is an indicator function. 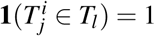 when 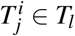, and 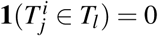 otherwise.

Furthermore, we can define the accuracy *Acc*_*ssp*_ of the *split sub-population* as follow.

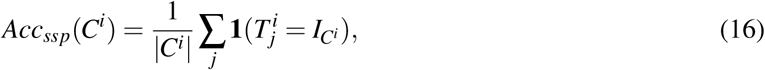

#### C.3 Annotation of a *common sub-population*

After finding the annotation 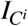 for each non-empty *split sub-population C*^*i*^ in *C*, we can check whether they have the same annotations. If all the non-empty *split sub-populations* in *C* are annotated to the some annotation, we consider the *common sub-population* as a *corresponding common sub-population* and use 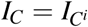 to denote the annotation for the *corresponding common sub-population*, where *i* is the index of a non-empty *split sub-population* in *C*. Otherwise, if the non-empty *split sub-populations* are annotated to different annotations, we consider the *common sub-population* as a *non-corresponding common sub-population* and set *I*_*C*_ = *NA*.

#### C.4 The ratio *R*_*ccsp*_ of the *corresponding common sub-population*

*R*_*ccsp*_ is the ratio of the number of the *corresponding common sub-populations* that identified by a method to the number of all identified *common sub-populations*. For example, if all *common sub-populations* are *corresponding common sub-populations, R*_*ccsp*_ = 100%; If all *common sub-populations* are *non-corresponding common sub-populations, R*_*ccsp*_ = 0%

Let ℭ = {ℭ_*I*_, ℭ_*c*_} be the output of a method, where ℭ_*I*_ is the set of all *individual sub-populations* and ℭ_*c*_ is the set of all *common sub-populations*. The ratio of the *corresponding common sub-population R*_*c*_ is defined as follow:

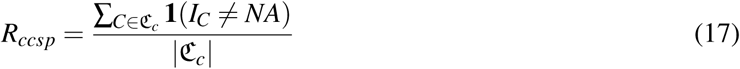

#### C.5 The ratio *R*_*ucct*_ of uncoveblack common cell types

*R*_*ucct*_ measures the percentage of the common cell types that can be uncoveblack. Let *T*_*c*_ be the set of ground truth labels that share by multiple data sets. We know *T*_*c*_ *∈ T, T* = {*T*_1_, *T*_2_, *…, T*_*L*_} and then *R*_*ucct*_ can be computed as

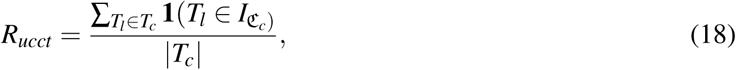

where 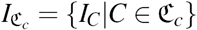.

#### C.6 The ratio *R*_*uict*_ of uncoveblack individual cell types

*R*_*uuct*_ measures the percentage of the individual cell types that can be uncoveblack. Let *T*_*I*_ be the set of ground truth labels that only appear in one data set. We know *T*_*I*_ *∈ T, T* = {*T*_1_, *T*_2_, *…, T*_*L*_} and then *R*_*uict*_ can be computed as

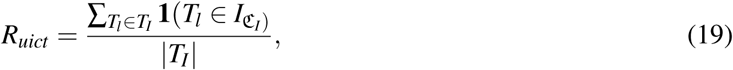

where 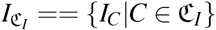.

#### C.7 The accuracy *Accc* of identified *common sub-populations*

Here we use *Acc*_*csp*_ to evaluate the purity of a *common sub-population*. Let *C* denote a *common sub-population* and 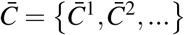 be the *common sub-population* where 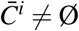;. The accuracy *Acc*_*csp*_ of the *common sub-population* of *C* can be computed as

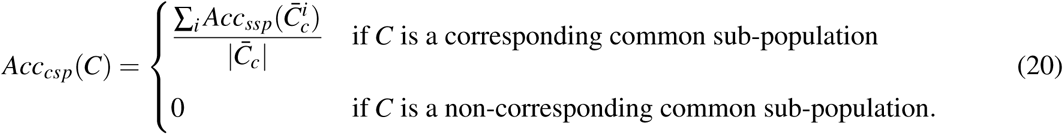

For the set ℭ of all *common sub-populations*, the accuracy of ℭ is

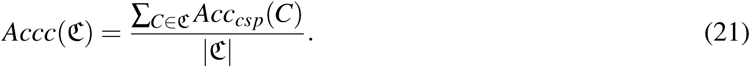

#### C.8 The accuracy *Acci* of identified *individual sub-populations*

The *Acc*_*isp*_ score for an *individual sub-population C*_*I*_ is

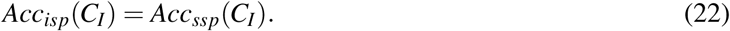

For the set ℭ_*I*_ of all *individual sub-populations*, the accuracy of ℭ_*I*_ is

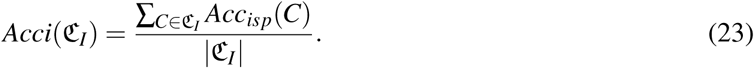

### D. Implementation details

#### D.1 Parameter settings

For human and mouse pancreatic single cell RNA-seq data sets, we set *λ* and *k* (the number of sub-populations), the only two parameters of PopCorn, to *λ* = 1 and *k* = 10, 11, 12, 13, 14, 15, respectively. We find that when we set *k* = 14, our PopCorn only generates 13 clusters, therefore, we use the results of *k* = 13 as our final results to compare with Seurat. For Seurat, we use the results they reported in [15] to compare with our PopCorn results because we use exact the same data sets. For Scanorama, we used the default parameters. And we use the Louvain algorithm to cluster the integrated data. We try the resolution parameters {0.2, 0.4, 0.6, 0.8, 1.0, 1.2} used by Louvain algorithm and pick the one yielded the best result.

For mouse kidney single cell data sets, we set *λ* = 3 and *k* = 19, cause when we set *k* = 20, PopCorn only generates 19 sub-populations. For Seurat, we use different number of HVGs (500, 1000, 1500) for the alignment method. For Scanorama, we use the default parameters. And for the sub-population identification method after Seurat alginment and Scanorama, we set the resolution parameters to {0.2, 0.4, 0.6, 0.8, 1.0, 1.2}. We try all combinations of the above parameter settings and show the best results, respectively.

#### D.2 Details results for human and mouse pancreatic data sets

Here we provide detailed comparison results (Fig. 4) of PopCorn, Seurat + Louvain, and Scanorama + Louvain for human and mouse single-cell RNA-seq pancreatic data sets. As shown in Fig. 4 and Table 1, PopCorn outperforms Seurat + Louvain and Scanorama + Louvain in terms of all metrics expect *A*_*cci*_, the accuracy of the identified individual cell populations. However, Scanorama + Louvain fails to find correspondence between beta cells in human and mouse (cluster # 12 and cluster # 14 in Fig. 4 C), which are 16.8% and 45.9% of the total cells in human and mouse data sets, respectively.

**Table 1:** Comparison on different metrics on human and mouse sinlge-cell RNA-seq data sets

| | $R_{ccsp}$ | $R_{ucct}$ | $R_{uict}$ | $A_{ccc}$ | $A_{cci}$ |
| --- | --- | --- | --- | --- | --- |
| PopCorn | **10/10** | **8/10** | **2/3** | **93.6%** | 90.9% |
| Seurat+Louvain | 8/10 | 7/10 | 0/3 | 71.4% | 39.3% |
| Scanorama+Louvain | 9/10 | 7/10 | **2/3** | 89.3% | **93.9%** |

**Fig. 4:**
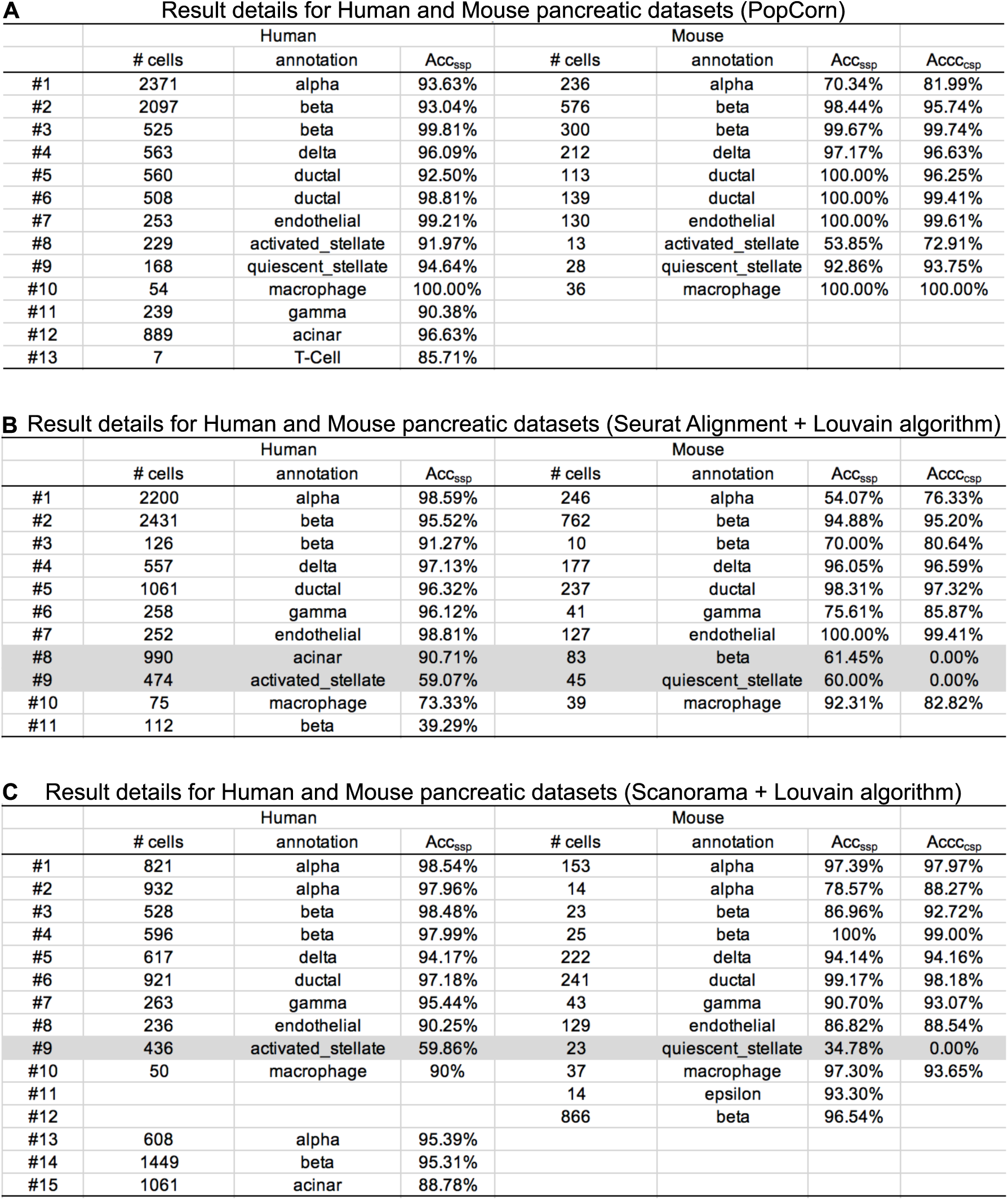
(A) Table for the detailed results for PopCorn on human and mouse pancreatic data sets. (B) Table for the detailed results for Seurat Alignment + Louvain algorithm on human and mouse pancreatic data sets. (C) Table for the detailed results for Scannorama + Louvain algorithm on human and mouse pancreatic data sets. Gray shade shows the *non-corresponding common sub-populations*.

#### D.3 Details results for mouse kidney data sets

Here we provide detailed comparison results (Fig. 5) of PopCorn, Seurat + Louvain, and Scanorama + Louvain for mouse kidney single-cell RNA-seq data sets. For this challenging data set, PopCorn significantly outperforms the rest in all metrics shown in Table 2.

**Table 2:** Comparison on different metrics on mouse kidney replicas data sets

| | $R_{ccsp}$ | $R_{ucct}$ | $A_{ccc}$ |
| --- | --- | --- | --- |
| PopCorn | **18/18** | **11/16** | **92.4%** |
| Seurat+Louvain | 14/19 | 6/16 | 76.8% |
| Scanorama+Louvain | 17/20 | 6/16 | 77.5% |

**Fig. 5:**
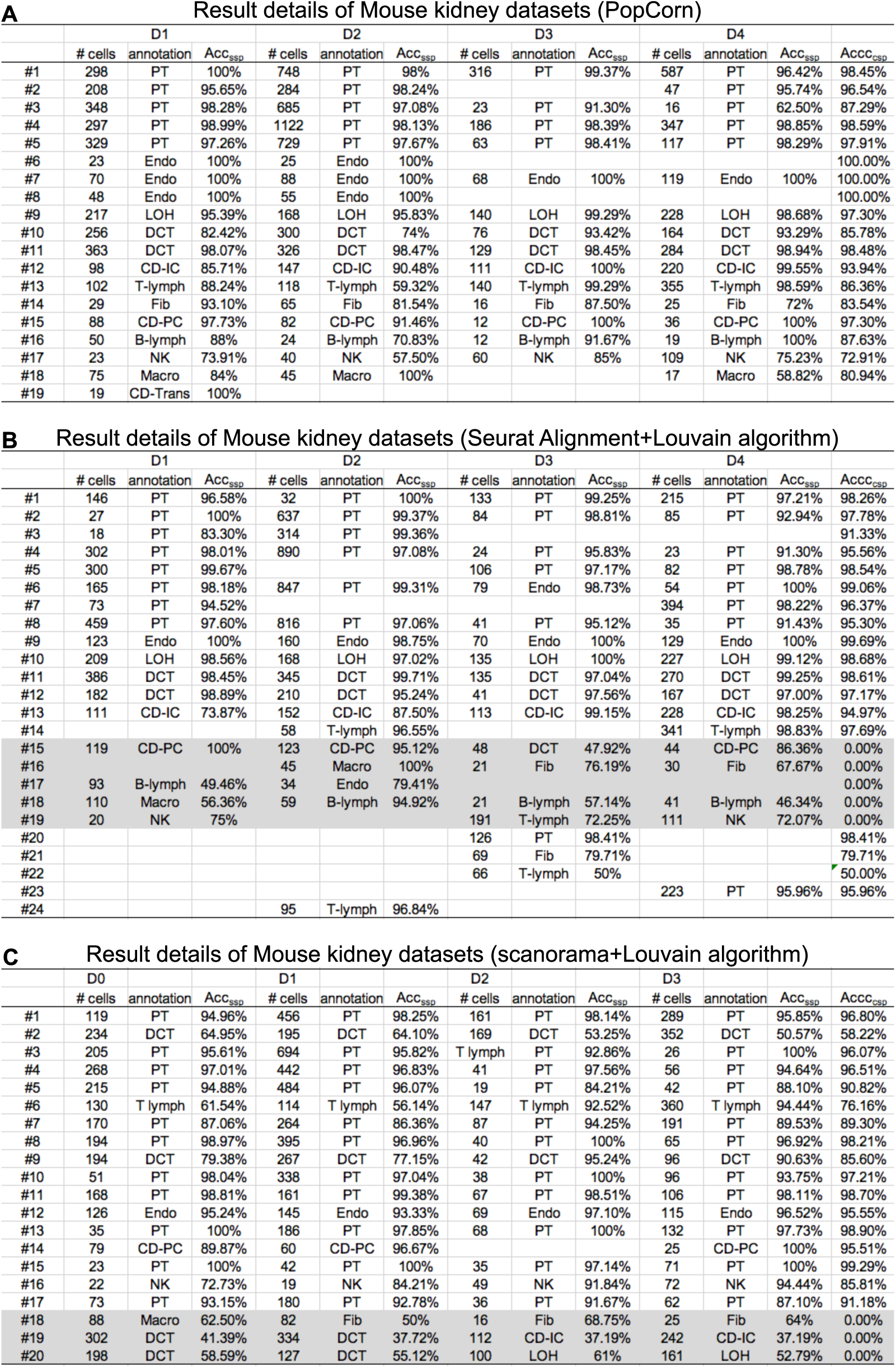
(A) Table for the detailed results for PopCorn on mouse kidney 4 replicas. (B) Table for the detailed results for Seurat on mouse kidney 4 replicas. Gray shade shows the *non-corresponding common sub-populations*.

